# U-shape short-range extrinsic connectivity organisation around the human central sulcus

**DOI:** 10.1101/2020.05.07.082800

**Authors:** Alexandre Pron, Christine Deruelle, Olivier Coulon

## Abstract

The central sulcus is probably one of the most studied folds in the human brain, owing to its clear relationship with primary sensory-motor functional areas. However, due to the difficulty of estimating the trajectories of the U-shape fibres from diffusion MRI, the short structural connectivity of this sulcus remains relatively unknown. In this context, we studied the spatial organization of these U-shape fibres along the central sulcus. Based on high quality diffusion MRI data of 100 right-handed subjects and state-of-the-art pre-processing pipeline, we first define a connectivity space that provide a comprehensive and continuous description of the short-range anatomical connectivity around the central sulcus at both the individual and group levels. We then infer the presence of five major U-shape fibre bundles at the group level in both hemispheres by applying unsupervised clustering in the connectivity space. We propose a quantitative investigation of their position and number of streamlines as a function of phenotypic traits such as sex and hemispheres and functional scores such as handedness and manual dexterity. Main findings of this study are twofold: a description of U-shape short-range connectivity along the central sulcus at group level and the evidence of a significant relationship between the position of three hand related U-shape fibre bundles and the handedness score of subjects.

## Introduction

The study of human brain connections terminations, i.e., the structural connectivity, is of great interest (Sporns et al. 2005), in particular, to broaden our knowledge of neurodevelopmental processes (e.g. (Dubois et al. 2014)), functional organization (e.g. (Saygin et al. 2011; Wendelken et al. 2017)) and brain diseases (e.g. (Griffa et al. 2013)). At the macroscopic scale, the connections of the white matter are organized into fibre bundles called tracts and qualified as commissural, projection and association tracts according to the location of their terminations (e.g. (Mandonnet et al. 2018)).

Short association tracts (Meynert 1885), also referred to as U-shape fibres or U-fibres, connect two cortical areas located in adjacent gyri, traveling in the superficial white matter, right beneath the sixth cortical layer (Schmahmann and Pandya 2006). These U-fibres, that account for about ninety percent of the intra-hemispheric connections (Schuz and Braitenberg 2002) are of most importance for anatomo-functional correspondence as a main and direct anatomical substrate of functional exchanges.

However, studies investigating their spatial distribution, variability or their functional roles across the human brain remain scarce (see e.g. (Guevara et al. 2017a) and references therein). This lack of knowledge may be due to the difficulty of reconstructing U-fibres based on diffusion-weighted magnetic resonance imaging (dMRI), the unique imaging modality that estimate the trajectories of white matter tracts, the streamlines, *in vivo*, in a non-invasive way. Indeed, U-fibres are short, curved, highly variable (e.g. (Dejerine and Déjerine-Klumpke 1895)) and located near the grey matter/white matter interface (GM/WM), a region affected by partial volume effect in dMRI. Superficial white matter also hosts complex fibre configurations potentially leading to erroneous streamlines reconstruction (e.g. (Reveley et al. 2015; Schilling et al. 2018)).

On location where a few studies of U-shape fibres have been produced is the central sulcus (CS), a deep, quite stable fold with respect to both topology and morphology of the brain (Ono et al. 1990), that splits apart two main functional regions, the primary motor and the primary sensory areas. For instance, Catani et al., (2012), identified U-fibres tracts coursing around the CS and confirmed it with *ex vivo* dissections, assessing the ability of dMRI and associated reconstruction methods to estimate U-fibres trajectories around this sulcus. However, despite a quite extensive knowledge of the CS with respect to cytoarchitectonic (e.g. (White 1997)), morphometry (e.g. (Cykowski et al. 2008; Sun et al. 2015)) and anatomo-functional correspondence (e.g. (Sun et al. 2016; Akselrod et al. 2017; Schellekens et al. 2018; Viganò et al. 2019; Germann et al. 2019)) studies focusing on the characterization of short association fibres around the human CS remain scarce.

A leftward asymmetry of the U-shape bundles volume was reported in (Catani et al. 2012) on 12 right-handed subjects. In (Rojkova et al. 2016), the methodology of (Catani et al. 2012) was applied to 47 subjects to assess the effects of age and education on the volume and on diffusion derived measures along the identified U-bundles of the CS. No significant effect of either age or education level was evidenced. Magro et al., (2012) found a significant leftward asymmetry of the U-fibres volume in left-handers that was correlated to handedness score, but no asymmetry was reported in right-handers. In (Román et al. 2017), a high-resolution template of the superficial white matter fibres was built from 74 subjects using both intra- and inter-subject clustering. This template was used to identify four reproducible U-shape bundles in the CS of 78 additional subjects. Contrary to (Catani et al. 2012; Magro et al. 2012), a significant rightward asymmetry was reported for two of the bundles located in the hand functional area. Lastly, Thompson and al., (2017) found a correlation between manual dexterity scores of both hands and microstructural features of the left hemisphere U-bundles within the hand but not the face area, suggesting a direct implication of these U-fibres in fine motor control.

Although providing worthwhile clarifications on the structural variability and on the functional role of the U-fibres of the CS, quoted studies reported inconsistent results preventing from drawing clear-cut conclusions. Differences between the results could be explained by either the heterogeneity of dMRI data quality, preprocessing algorithms, local models and tract selection strategies used.

Typical dMRI data used in the above-mentioned studies are mostly of lower spatial (e.g. 2 mm) and angular resolution (e.g. (64 directions, b=1500 s.mm^−2^)) than those currently available in large databases such as the Human Connectome Project (HCP) Young Adult dataset^1^ (e.g. (1.25 mm and 3 times 90 directions, b=1000, 2000, 3000 s.mm^−2^)). In addition, progress in the dMRI data preprocessing, local modelling and in the algorithms involved in streamlines reconstruction make them better handle partial volume effect and complex crossing that occur in the cortex vicinity. There is thus a clear need to provide a complete representation of the short structural connectivity of the CS as well as a characterization of the U-shape bundles of the central area, relying on large sample of high quality dMRI data and appropriate processing methods.

The present study has two objectives. Firstly, we aim at providing a more exhaustive and accurate estimation of U-fibres of the CS from HCP dMRI data of one-hundred subjects processed with state-of-the-art modelling, tractography and clustering algorithms. Secondly, we aim at extracting consistent U-bundles of the CS across subjects and characterizing them with respect to lateralization, sex, handedness and motor scores.

## Material and Methods

### Participants

One-hundred right-handed (manual laterality score derived from the Edinburgh Handedness Inventory (Oldfield 1971) greater than or equal to 50), non-twin subjects (50 males, 50 females), aged between 22 and 36 years old (*M* = 28.82, *SD* = 4.04), were selected from the publicly available HCP Young Adult dataset (Van Essen et al. 2013). Male and female subgroups had similar age distribution (non-parametric Kolmogorov-Smirnov test, *D* = 0.12, *p* = 0.84).

### MRI data acquisition

MRI data were acquired by the HCP using a modified version of Siemens Skyra 3 T scanner (Siemens, Erlangen, Germany) with a maximum gradient strength of 100 mT. m^−1^, slew rate of 91 T. m^−1^. s^−1^ and a 32-channel head coil.

T1 weighted (T1w) MRI images were acquired using three-dimensional Magnetization Prepared Rapid Gradient Echo (3D-MPRAGE) sequence (repetition time (TR) / echo time (TE) = 2400/2.14 ms, flip angle = 8 °, resolution = 0.7 mm isotropic, field of view (FOV) = 224 × 224 mm^2^).

T2 weighted (T2w) MRI scans were acquired using a variable flip angle turbo spin-echo sequence (TR/TE = 3200/565 ms, resolution = 0.7 mm isotropic, same FOV than for T1w images).

dMRI scans were acquired using a spin-echo, echo planar imaging sequence consisting of three shells (*b* = 1000, 2000 and 3000 s. mm^−2^) of 90 volumes each and 36 interleaved non diffusion weighted volumes (*b* = 0 s. mm^−2^) (TR/TE = 5520/89.5 ms, resolution = 1.25 mm isotropic, FOV = 210 × 180 mm^2^, 111 axial slices, multiband factor = 3, partial Fourier = 6/8, echo spacing = 0.78 ms). Gradients directions were sampled over the entire sphere, using the electrostatic repulsion method (Caruyer et al. 2013). The whole diffusion sequence was repeated twice with reversed phase encoding (left to right, right to left).

Further information about MRI data acquisition can be found in (Glasser et al. 2013). All selected data have been through control quality carried out by the HCP (Marcus et al. 2013).

### Structural MRI data processing

Undistorted and *B*_1_ bias corrected T1w and T2w scans in subject native space as provided by the *PreFreeSurfer* pipeline version 3.1.3 (https://github.com/Washington-University/HCPpipelines) (Glasser et al. 2013), were processed with FreeSurfer 6.0.0 (https://surfer.nmr.mgh.harvard.edu) (Fischl 2012), to benefit from the submillimetre resolution full support brought by this version. GM/WM and pial triangular meshes were reconstructed from the FreeSurfer tissue segmentation relying on the Morphologist pipeline (Fischer et al. 2012) of the BrainVISA 4.6.1 suite (http://brainvisa.info) (Geffroy et al. 2011). A visual quality control was carried out on each generated GM/WM mesh to ensure that it was free of any significant topological or geometrical defects.

Surface maps of mean curvature, geodesic depth and depth potential function (DPF) (Boucher et al. 2009) were obtained resorting to the BrainVISA Cortical Surface toolbox (http://brainvisa.info/web/cortical_surface.html) with default parameters.

In order to further extract short association fibres around the CS, we need to extract cortical landmarks related to this sulcus. Specifically, the crests of the pre-central gyrus (PreCG), the post-central gyrus (PoCG), and the CS fundus lines were semi-automatically delineated onto each GM/WM mesh relying on the SurfPaint module (Le Troter et al. 2011) of the Anatomist software (Rivière et al. 2011). This software can extract gyrus crest lines (resp. sulcal fundus lines) as geodesic lines between two extremities, with these geodesic lines minimizing (resp. maximizing) the DPF along their path (see e.g. (Le Troter et al. 2012)). Extremities of each of the three lines were positioned manually. The dorsal endpoint of the three lines was set at the *apex* of the characteristic notch made by the CS on the medial face of the GM/WM mesh. The ventral endpoint was set on the crest of the sub-central gyrus (SubCG). The procedure followed to locate the SubCG and to position the endpoints of the CS is detailed and illustrated in the Supplementary Material. Regarding the PreCG and the PoCG crests lines extraction, four manually specified controls points were added to differentiate between the two gyri and to ensure an accurate delineation in bending areas. The placement of these control points is described and illustrated in the Supplementary Material.

The position of the *pli de passage fronto-pariétal moyen* (PPFM) (Broca 1888), an *annectant gyrus,* located in the midpart of the CS, which is a surface landmark of the primary motor area of the hand (Boling and Reutens 1999), was drawn on each CS fundus line and projected onto each delineated gyral crest. The drawing procedure followed is detailed and illustrated in the Supplementary Material.

### dMRI data processing

dMRI scans pre-processed by the HCP, i.e., corrected for subject movement, susceptibility induced artefacts, eddy-current induced distortions and gradients non linearities (Jenkinson et al. 2012; Glasser et al. 2013), were used to build whole brain tractograms with Mrtrix3 (http://www.mrtrix.org) (Tournier et al. 2019). Pre-processed scans were first corrected for non-uniform intensity adopting the ANTS (https://github.com/ANTsX/ANTs) N4 bias correction algorithm (Tustison et al. 2010). For each subject, a multiple shell multiple tissue (MSMT) (cerebrospinal fluid, grey matter, white matter) response function was derived from the FreeSurfer tissue segmentation relying on the *dwi2response* command with default parameters. The resulting response function was used to fit a constrained MSMT spherical deconvolution model (Jeurissen et al. 2014) on the brain diffusion signal. Whole brain probabilistic tractography (Tournier et al. 2010) was performed with the *tckgen* command (algorithm = iFOD2, step = 0.625 mm, angle = 45°, nb_streamlines = 5 × 10^6^, minlength = 2.50 mm, maxlength = 300 mm) with seeding from the GM/WM interface volume and anatomical constraints (e.g. endings in the grey matter, see (Smith et al. 2012)) stemming from the FreeSurfer tissue segmentation.

The generated tractograms were filtered within the *Convex Optimization Modeling for Microstructure Informed Tractography* (COMMIT) framework (https://github.com/daducci/COMMIT) (Daducci et al. 2013, 2015) in order to remove spurious or overrepresented streamlines with respect to the dMRI signal. Stick-Zeppelin-Ball (Panagiotaki et al. 2012) with default diffusivity parameters (parallel diffusivity = 1.7 × 10^−3^ mm^2^.s^−1^, intracellular fraction = 0.7, isotropic diffusivities = 1.7 × 10^−3^ and 3.0 × 10^−3^ mm^2^.s^−1^) was selected as a forward model (Daducci et al. 2015). The resulting tractograms contained an average of one million streamlines, which corresponds to a reduction of eighty percent.

The association streamlines, i.e., connecting ipsilateral cortical territories, were extracted from the filtered tractogram, using their endpoints and a binary mask of each hemisphere cortical grey matter. These streamlines were further filtered with respect to the GM/WM mesh as follows. The exact signed distance from the streamlines points and the GM/WM mesh was computed relying on trimesh (https://github.com/mikedh/trimesh)(Dawson-Haggerty et al. 2019). Each segment composing a streamline was classified into intersection, intracortical (grey matter) or subcortical (white matter) using the values of the signed distance at the mesh of its endpoints. Streamlines with more than two intersections with the mesh were excluded. Regarding streamlines with at most two intersections, only the intersections and the subcortical components were retained, with the latter having a length greater than the minimum value of 2.50 millimetres.

### CS U-shape streamlines extraction

From the resulting tractograms we then extracted short U-shape association fibres around the CS. For this, two regions of surface interest (ROI) corresponding to the PreCG and the PoCG were generated. For each of the two gyri, we used the corresponding crest line and the CS fundus line. The corresponding ROI was defined as the GM/WM mesh vertices located closer to the crest line than the local geodesic distance between the crest line and the CS fundus line.

Streamlines terminations were projected onto the GM/WM mesh points by minimizing the Euclidean distance. U-shape streamlines of the CS were then extracted as streamlines with their two terminations projected on the two ROIs (PreCG and PoCG). The top one percent of the longest U-streamlines in each of the CS were filtered out in order to remove obvious outliers such as looping streamlines. An example of the resulting set of streamlines is shown in **Fig. *2***.

**Fig. 1.**
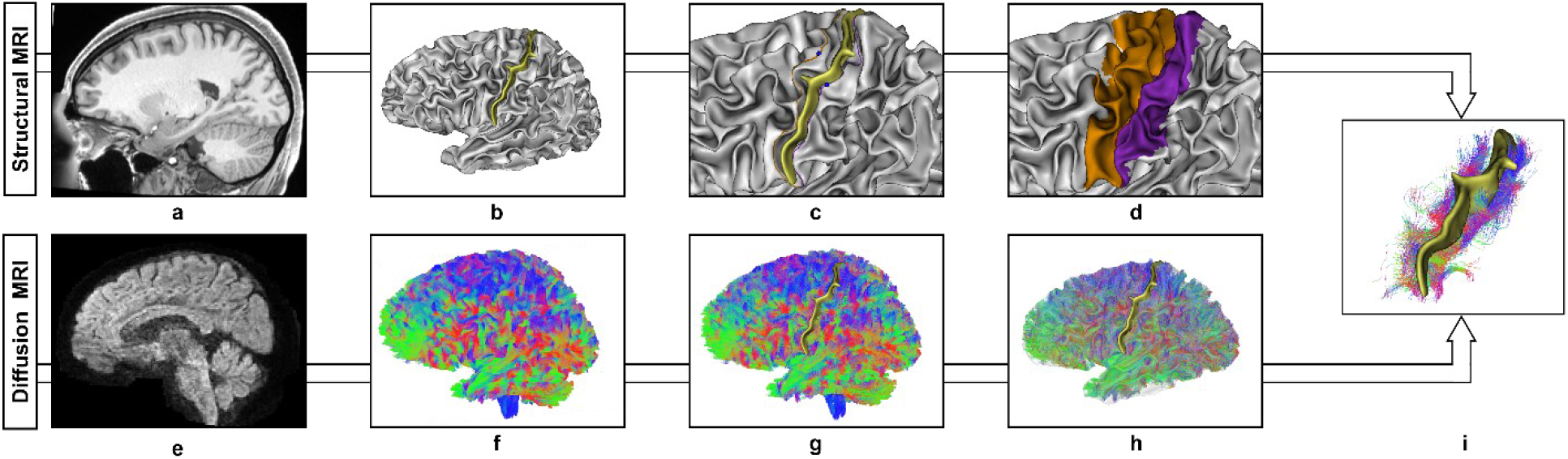
Central sulcus U-shape streamlines reconstruction pipeline illustrated on a randomly selected subject (id:114621) **a** T1w scan pre-processed by the HCP **b** Left grey matter/white matter (GM/WM) interface triangular mesh generated from the FreeSurfer 6.0.0 tissue segmentation and the associated central sulcus represented by a yellow ribbon. **c** Pre-central (orange) and post-central (purple) gyral crest delineated onto the GM/WM mesh. The pli de passage fronto parietal moyen along each crest is represented by blue spheres. **d** Pre-central (orange) and post-central (purple) surface area generated from gyral crests and central sulcus fundus line. **e** dMRI scan pre-processed by the HCP. **f** Whole brain probabilistic and anatomically constrained tractogram (5 million streamlines) color-coded by principal streamline direction (left-right: red, antero-posterior: green, top-bottom: blue). **g** COMMIT filtered tractogram (∼ 1 million streamlines). **h** Association streamlines of the left hemisphere filtered with respect to the GM/WM mesh. **i** U-shape streamlines of the left central sulcus, color-coded by local (segment) streamline directions (left-right: red, antero-posterior: green, top-bottom: blue).

**Fig. 2.**
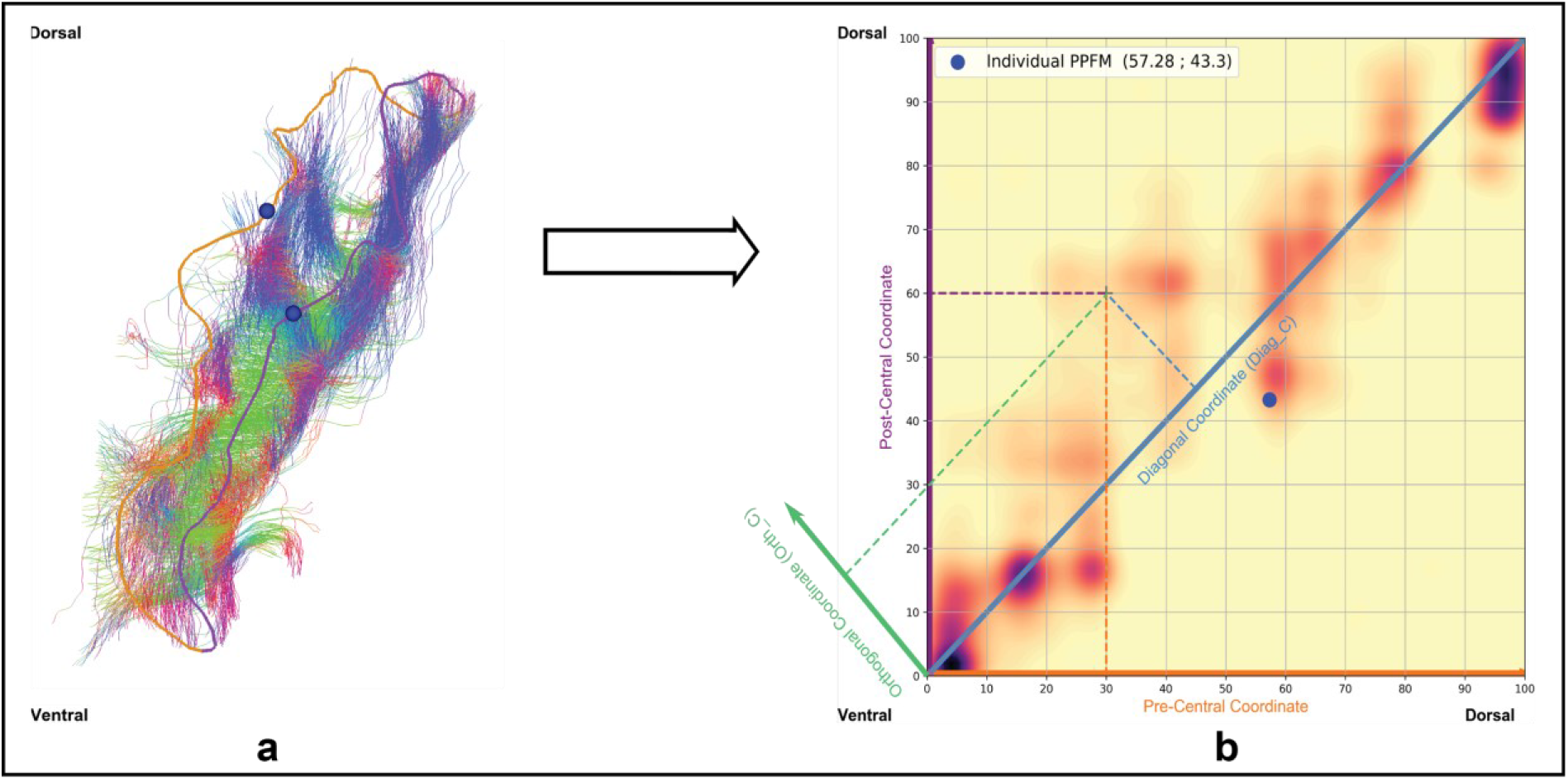
Individual connectivity space construction illustrated on one randomly chosen subject. **a** U-shape streamline terminations of the central sulcus are associated with a coordinate along both the pre-central(orange) and post-central (purple) crest lines. **b** These two lines being isometrically parameterized from their ventral (0) to their dorsal (100) extremity, U-shape streamlines structural connectivity can be represented in a continuous bidimensional space [0,100] × [0,100]. The continuous profile is obtained by interpolation by Gaussian kernel of scalar covariance matrix and standard deviation *σ* = 5. Anatomical information such as pli de passage fronto parietal median position can be added (blue dot). Other coordinates system such as the (Diagonal (dodger blue), Orthogonal (green)) coordinate system can be used

### Short structural connectivity profiles of the CS

We relied on the connectivity space framework first introduced in (Pron et al. 2018) to study the structural connectivity of U-fibres of the CS: a continuous space, *the connectivity space,* is defined in which each U-streamline is represented by a point whose coordinates are a function of the position of its endpoints along the adjacent gyral crests lines. The structural connectivity of the U-shape fibres around the CS is therefore represented by a set of points, the *connectivity profile*, in this bidimensional, continuous space.

We isometrically parameterized the PreCG and PoCG crest lines, in an intrinsic manner, as proposed in (Coulon et al. 2015), from the ventral (coordinate 0) to the dorsal end (coordinate 100). For a given streamline, its pre-central (resp. post-central) endpoint was projected onto the PreCG (resp. PoCG) line by geodesic distance minimization and the resulting coordinate is used. The streamline is then associated to two coordinates that define its position in the [0; 100] × [0; 100] connectivity space (see **Fig. *2***).

Assuming that the gyral crest lines are homologous between subjects and hemispheres, the normalized [0; 100] parameterization also guarantees a homology between connectivity spaces of different subjects (and different hemispheres). Therefore, connectivity profiles of different subjects can be gathered in the same space to provide a *group connectivity profile*. In order to improve the inter-individual correspondence, the individual positions of the PPFM were aligned across subjects on the mean groupwise PPFM position (left and right hemispheres combined) using piecewise affine transform as described in (Coulon et al. 2011).

In addition, we introduced a new coordinate system (Diag_C, Orth_C) of the connectivity space in order to provide anatomically interpretable descriptions of the spatial organisation of the U-fibres terminations of the CS (see **Fig. *2***). The diagonal coordinate (Diag_C) corresponds to the mean position of the streamline along the dorso-ventral axis of the CS. The orthogonal coordinate (Orth_C) quantifies a torsion with respect to this axis. In the case of a positive or negative Orth_C coordinate, the torsion is described as positive or negative.

### Inference of U-fibres bundles

Once the group connectivity profiles are built from a large number of streamlines represented in the group connectivity space, we want to extract bundles. To do so, group connectivity profiles of the left and the right hemisphere were clustered separately using the scikit-learn implementation (Pedregosa et al. 2011) of the *Density Based Spatial Clustering of Applications with Noise* (DBSCAN) algorithm (Ester et al. 1996) with (ϵ = 3, min_sample= 12000). The ϵ parameter value was determined following the heuristic proposed in (Ester et al. 1996). The min_sample parameter value was set through grid search on the domain *I* = [500,100000] with step 100 as the first value that produced clusters with smooth borders.

Three clusters were obtained bilaterally (**Fig. *4***): one in the ventral extremity of the space, one in the dorsal extremity of the space and a last one in the central part of the space. Visual inspection of the group connectivity profiles showed that for both hemispheres the central cluster contained three density peaks corresponding to three sub-clusters It was therefore sub-divided into three clusters using scikit-learn gaussian mixture model with default parameters. Inter-hemispheric correspondence between the bundles was achieved by sorting them with respect to the precentral coordinate of their centroids in the group space. Bundles were then numbered from the ventral (1) to the dorsal (5) extremity of the CS (**Fig. *4***).

**Fig. 3.**
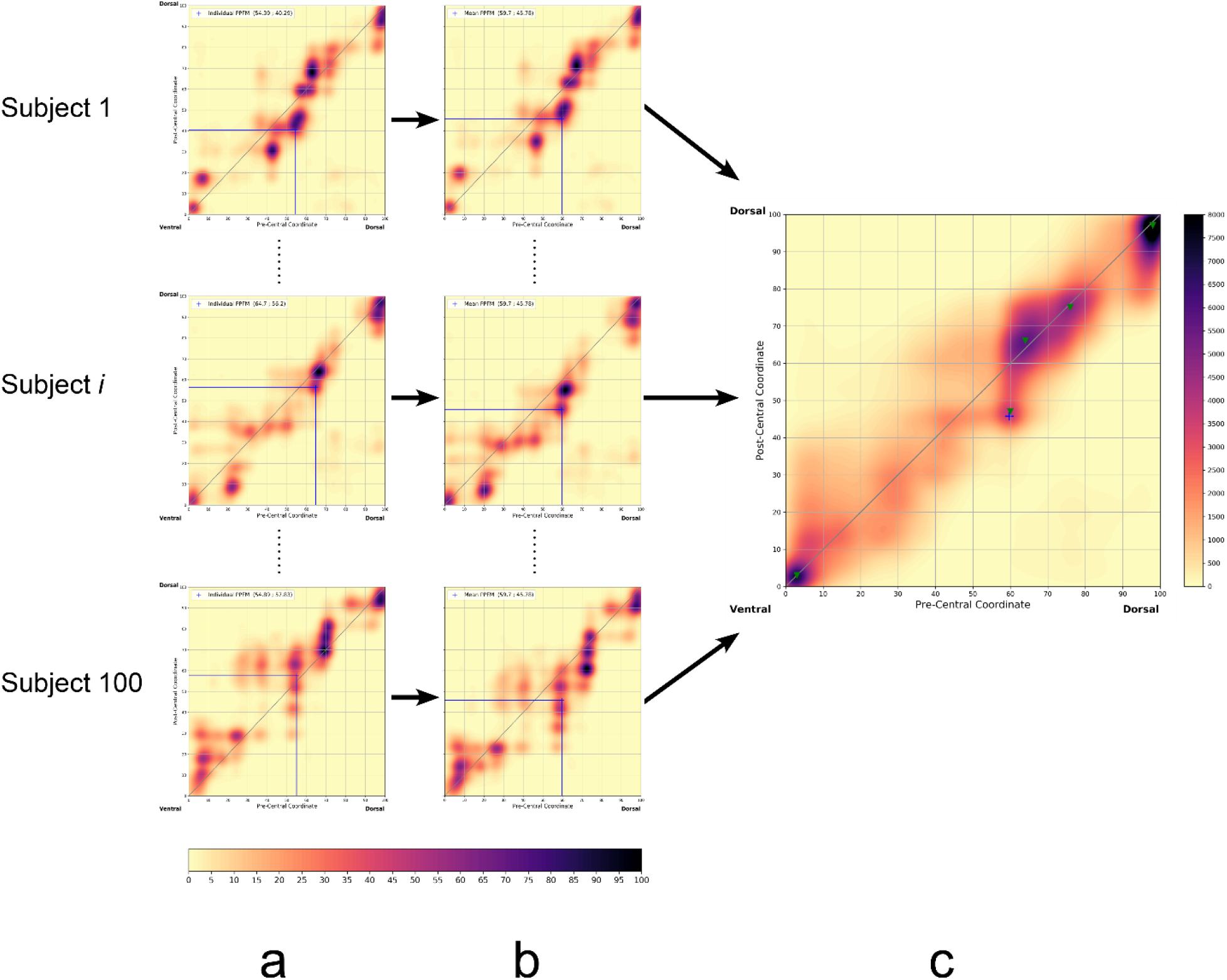
Group connectivity profile (**c**) construction from the 100 raw individual profiles (**a**), illustrated on left hemisphere. Individual PPFM positions (**a**, blue crosses) are first aligned onto the mean groupwise PPFM position (**b**, blue cross) by piecewise affine transform to improve anatomical correspondence across subjects. The group connectivity profile is the sum of the 100 aligned individual profiles ((**a**, **b**), common individual colorbar, (**c**) group colorbar).

**Fig. 4.**
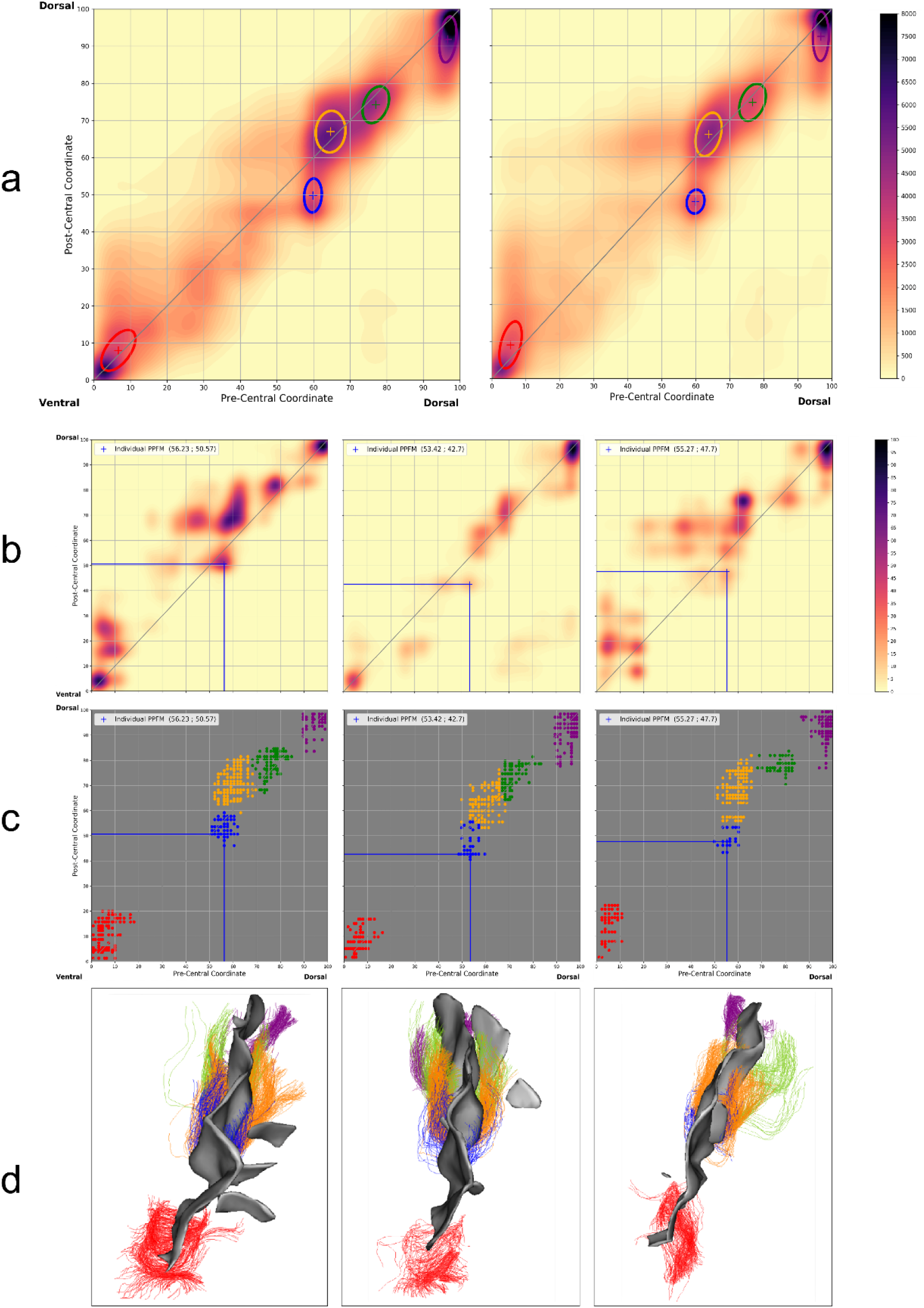
Extraction of U-shape fibre bundles of the central sulcus by classifying associated group connectivity profiles. The bundles obtained are represented at group level (**a**) by their centroid (crosses) and their covariance matrix (ellipse): 1 (red), 2 (blue), 3 (yellow), 4 (green), 5 (purple). These bundles are transported in the individual native connectivity spaces as shown in three randomly selected hemispheres ((**b**) individual connectivity profiles, (**c**) individual bundles). These bundles can also be viewed in three dimensions (**d**).

The five resulting clusters were backpropagated to each individual space and centroids were obtained by averaging over resulting points by cluster as shown in **Fig. *4***.

### Statistical analyses

Statistical analyses were performed with the Jamovi software (https://www.jamovi.org) and the statsmodels 0.9.0 Python module (Croxson 2005).

The handedness score (values between 50 and 100 in our sample) was recoded as a two level (Low: [50,85], n=57; High:]85,100], n=53) categorical variable dividing the sample into strongly (High) and weakly (Low) right-handers.

Streamlines count was performed for each cluster of both hemispheres. The volume of a bundle was defined as the volume in mm^3^ of the voxels containing at least one streamline of the bundle. Normalized asymmetry coefficients *AC* for streamline count and bundle volume were calculated from the formula: *AC* = 2 × (R - L) / (R + L) where R and L refer to a same quantity measured in right and left hemisphere, respectively. Negative value indicates a leftward asymmetry, zero value the absence of asymmetry.

A repeated measures ANOVA was used to evaluate the interaction between hemisphere (Left hemisphere, Right hemisphere), handedness category (Low and High right-handers), sex (Male, Female) and the position of the clusters’ centroid in the individual spaces. Position was determined relying on the (Diag_C, Orth_C) coordinates system and each coordinate was processed separately.

An ANOVA was used to assess the interaction between handedness (Low and High right-handers), sex (Male Female) and the streamline count asymmetry by cluster. Effect size of the ANOVAs was calculated using partial eta-squared 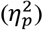 and results were reported using type 3 sum of squares.

Given the significant correlations reported between a motor score and volume of the bundles in the left CS hand functional area by (Thompson et al. 2017), a linear model was used to investigate the association between the streamline count of clusters 2,3,4 in the left hemisphere and dexterity score adjusted for age and covarying for sex. Dexterity score provided by the HCP is the Nine Holes Peg Test (Mathiowetz et al. 1985) score in the dominant hand that measures finger dexterity. The score is normed using the age appropriate band of Toolbox Norming sample (bands of ages 18-29 or 30-35); a score of 100 indicates performance at national average.

A similar model was used to assess the association between the streamline count inter-hemispheric asymmetry score of clusters 2,3,4 and a strength score, covarying for sex. Strength score provided by the HCP represents the relative force generated using the dominant hand. The strength score is normed with respect to the same band age previously described; a score of 100 indicates performance at national average. Further information about the Dexterity and Strength scores is available in the HCP dictionary.

All statistics reported used a level of significance, *α*, fixed to 0.05. Shapiro-Wilks tests were used to assess normality of the distributions (e.g. asymmetries, residuals of ANOVAs and linear models). ANOVAs homogeneity of variance hypothesis was assessed using the Levene test. Significance of the asymmetries was evaluated using either one sample t test or Wilcoxon-Mann-Whitney test depending on the result of normality test. Post-hoc comparisons of ANOVAs and repeated measures ANOVA were performed with p-values corrected for multiple comparisons using the Bonferroni method. Results of the post-hoc tests were reported in the form *t*(*dof*) =., *p* =., *n*.) where *n* stands for the number of tests performed. A whole number of degrees of freedom indicates that the Student test was used while a decimal number of degrees of freedom (*dof*) indicates that the Welch test was used.

## Results

Individual connectivity profiles of both hemispheres for the 100 subjects were computed, as well as group connectivity profiles.

### Qualitative description of left and right hemisphere group connectivity profiles

A continuous representation of the group profiles of both hemispheres group profiles shows a high streamline density along the diagonal of the space (see **Fig. *5***). This means an Orth_C coordinate close to 0, which indicates streamlines mostly orthogonal to the main axis of the sulcus, connecting homologous areas facing each other on the PreCG and on the PoCG. As presented before, three main high-density areas are visible in the ventral, the central and the dorsal parts of the group connectivity space of both hemispheres. Regarding the central part, three local maxima of the streamline’s density were present in both left and right hemispheres.

**Fig. 5.**
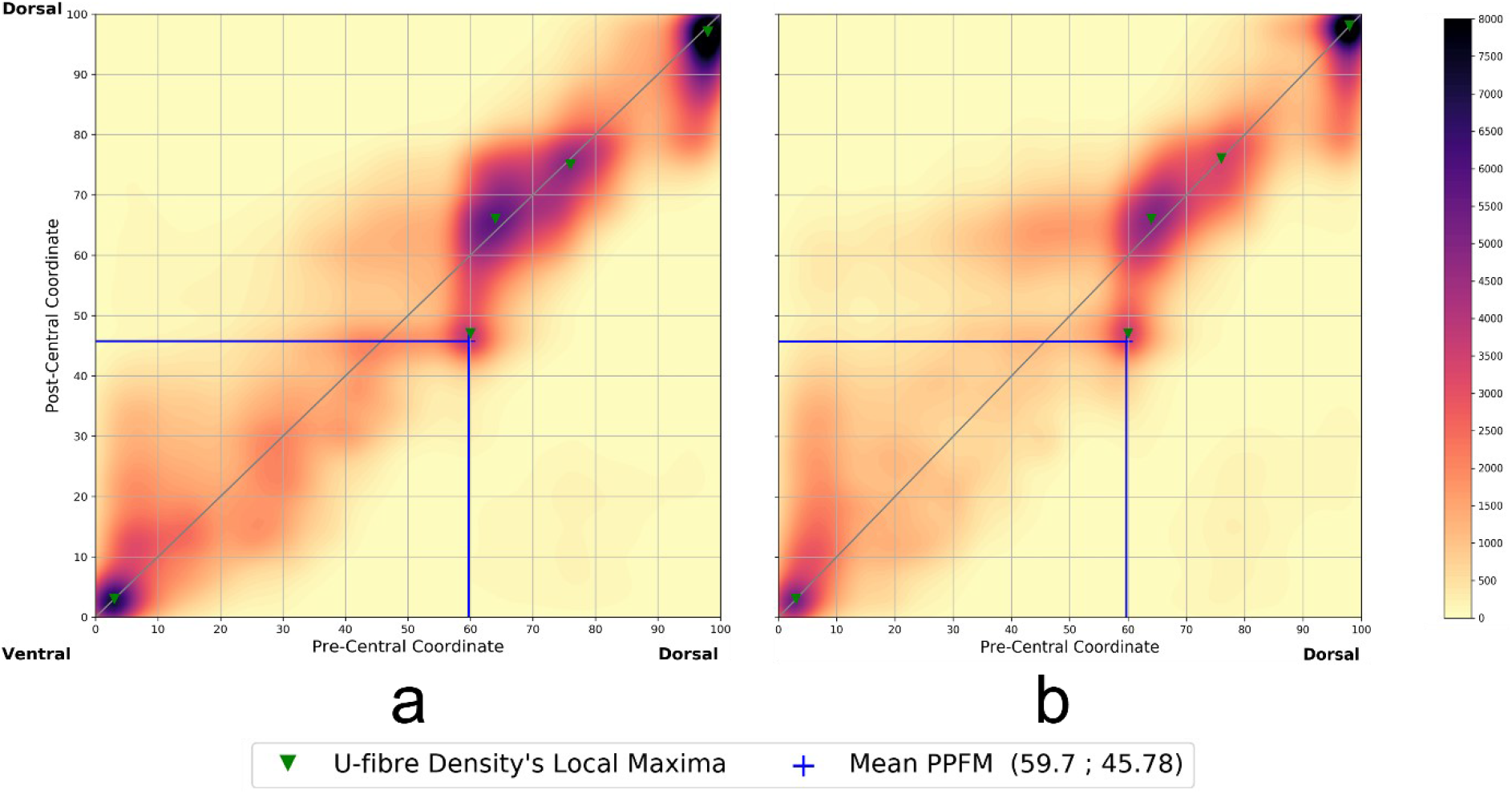
Connectivity profiles of the left (a) and right (b) central sulcus constructed from the profiles of the 100 subjects in the study showing concentration of streamlines along the diagonal with similar patterns between hemispheres. The continuous profile is obtained by interpolation by Gaussian kernel of scalar covariance matrix and standard deviation *σ* = 5 and depends on the number of streamlines. Local connectivity profile maxima (green crosses) are detected on a 2-neighbourhood and with a value greater than 20 percent of the value of the global maximum per profile

**Fig. 6.**
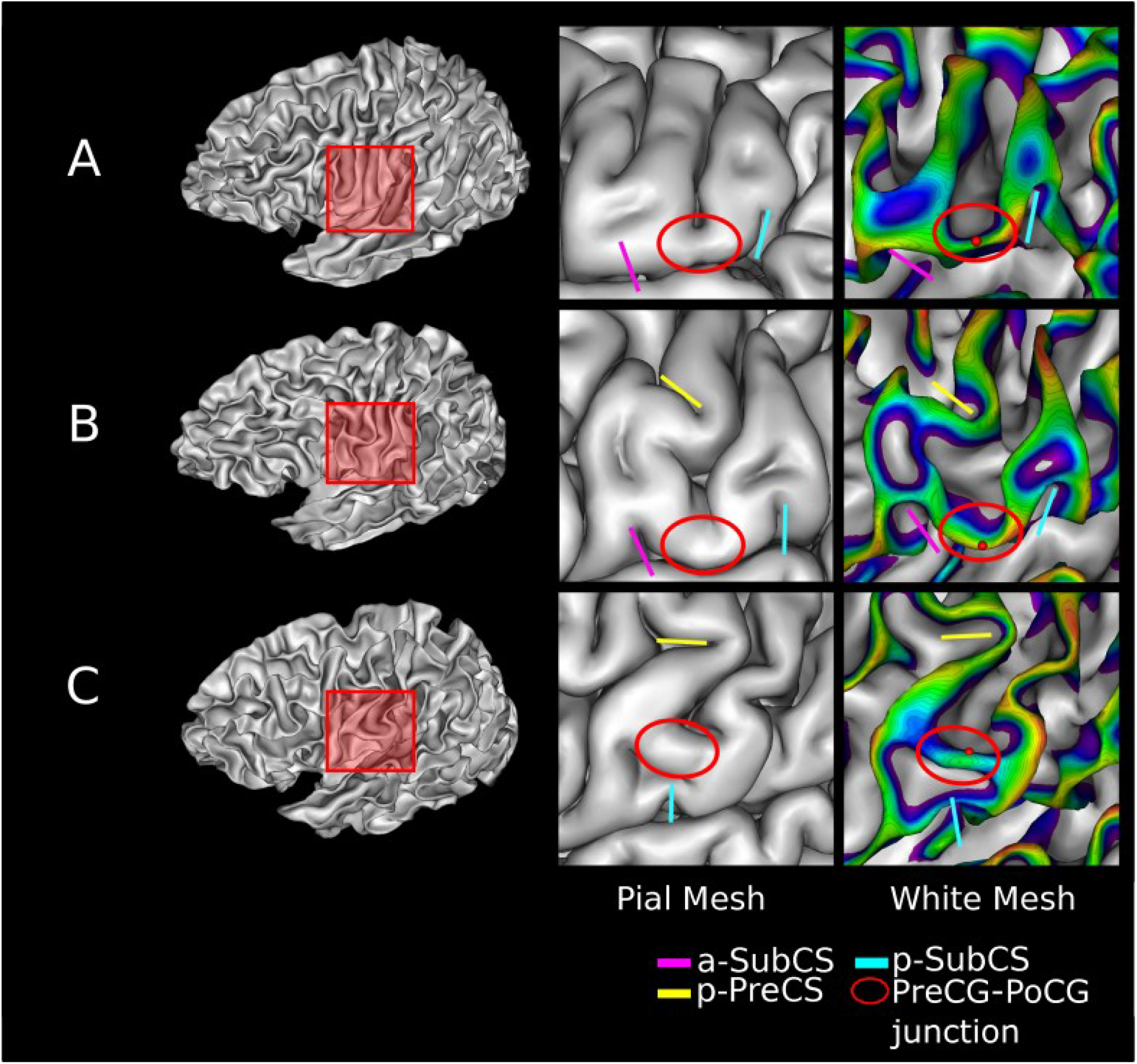
Positioning of the ventral end of the CS (red sphere) on three GM/WM meshes with different morphological configurations. The junction between the pre-central gyrus and the post-central gyrus (red ellipse) is located on the pial mesh (middle column) using adjacent sulci (anterior and posterior subcentral sulcus). The posterior branch of the precentral sulcus (p-precs) was also represented when visible. The sphere is located on the crest of the junction and minimizes the distance to the zero-level set of the DPF

**Fig. 7.**
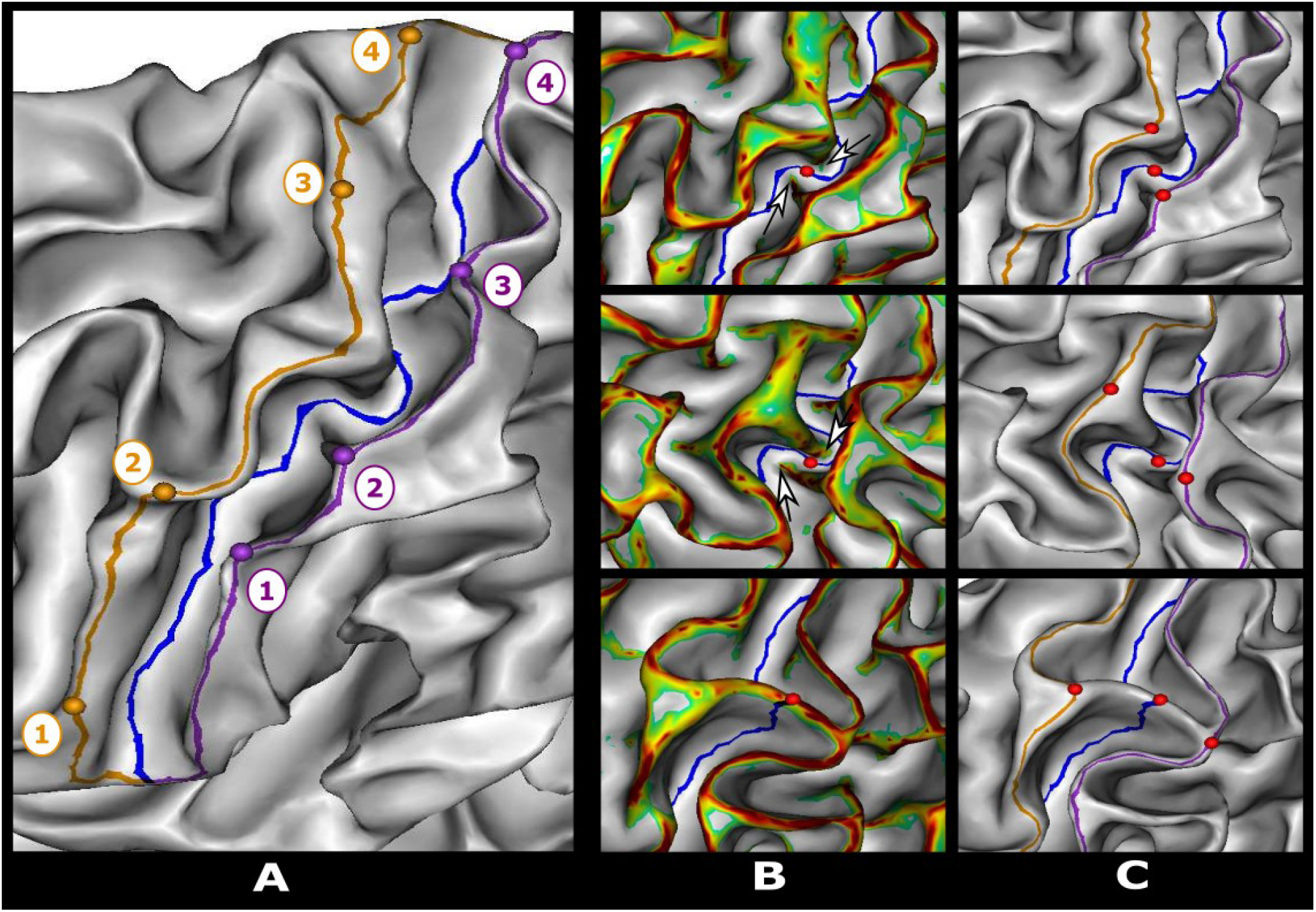
Left: semi-automatic delineation of the central sulcus fundus line (blue) and adjacent gyri ridge lines (orange pre-central gyrus, purple post-central gyrus) and their associated control points numbered in ascending order from the ventral end to the dorsal end of the central sulcus. Right: Manual positioning of the intersection of the intersection of the pli de passage fronto pariétal moyen (PPFM) with the bottom line of the central sulcus (red sphere) from the gyral protuberances (white arrows) marked on a thresholded mean curvature map. The position of the PPFM is then projected onto the ridge lines by minimizing the geodesic distance weighted by the mean curvature

Five U-bundles were obtained on the left and five bundles in the right hemisphere by clustering streamlines points in the group space. As visible on Figure 1, bundle 2 is located at the level of the mean position of the PPFM, below the diagonal of the space as seen in Figure 1, which denotes a negative Orth_C coordinate and thus a negative torsion with respect to the ventro-dorsal axis. Bundles 3 and 4 are just dorsal to the bundle 2, and thus to the PPFM, indicating that these 3 bundles correspond to the position of the primary hand area. Bundle 1 is the most ventral one, corresponding to the orofacial functional area whereas bundle 5 is the most dorsal one, corresponding to the foot/leg functional area. In the following, we quantified the effect of gender, handedness category and hemisphere on the position of the centroids and the number of streamlines in each of the extracted fibre bundles.

### Effect of hemisphere, handedness, and sex on bundle’s centroids position

An analysis of variance with repeated measures was carried out on the diagonal (*Diag_C*) and on the orthogonal (*Orth_C*) coordinates of the five centroids in individual connectivity spaces to assess the effect of hemisphere, sex and handedness category (Low or High right-handers). Results by centroid and coordinate are described below.

- Centroid 1:

– Diagonal: A significant main effect of the hemisphere (*F*(1,96) = 7.618, *p* = 0.007, 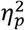 = 0.074) was reported so that centroid was more dorsal in the left than in the right hemisphere. A significant interaction between handedness category and sex (*F*(1,96) = 5.376, *p* = 0.023, 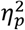 = 0.053) was shown but none of the post-hoc tests was statistically significant even before Bonferroni correction for multiple comparisons (all *p* ≥ 0.05).
– Orthogonal: Significant main effects of the hemisphere (*F*(1,96) = 11.766, *p* < 0.001, 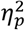 = 0.109) and of the handedness category (*F*(1,96) = 10.71, *p* = 0.03, 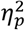 = 0.048) were reported so that absolute orthogonal coordinate was larger in the right than in the left hemisphere and also larger in the Low than in the High right-handers.
- Centroid 2:

– Diagonal: A significant interaction between hemisphere and handedness category (*F*(1,96) = 10.824, *p* = 0.001, 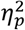 = 0.101) was observed. Centroid was more dorsal in the left than in the right hemisphere in the High right-handers (*t*(96) = 5.28, *p* < 0.001, *n* = 4) but not in the Low right-handers (*t*(96) = 1.09, *p* = 1.00, *n* = 4). Besides, centroid was more dorsal in the High right-handers compared to the Low right-handers, in the left (*t*(185.8) = 2.81, *p* = 0.02, *n* = 4) but not in the right hemisphere (*t*(185.8) = −1.39, *p* = 0.66, *n* = 4).
– Orthogonal: A significant main effect of the hemisphere (*F*(1,96) = 64.8992, *p* < 0.001, 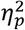 = 0.403) was shown, so that the absolute value of the orthogonal coordinate was larger in the left than in the right hemisphere. A significant interaction between sex and handedness category (*F*(1,96) = 10.71, *p* = 0.036, 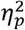 = 0.045) was also reported: Low right-handers had a larger absolute value of the orthogonal coordinate than the High right-handers in the male (*t*(96) = −2.741, *p* = 0.028, *n* = 4) but not in the female subgroup (*t*(96) = 0.192, *p* = 1, *n* = 4). No significant difference was shown between males and females neither in the Low right-handers (*t*(96) = −1.876, *p* = 0.256, *n* = 4) nor in the High right-handers (*t*(96) = 1.184, *p* = 1, *n* = 4).
- Centroid 3:

– Diagonal: A significant interaction between the hemisphere and the handedness category factors (*F*(1,96) = 16.965, *p* < 0.001, 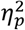= 0.150) was found. Coordinate was more dorsal in the left than in the right hemisphere in the High right-handers (*t*(96) = 5.003, *p* < 0.001, *n* = 4) but not in the Low right-handers (*t*(96) = 0.500, *p* = 1, *n* = 4). In addition coordinate was more dorsal in the High right-handers than in the Low right-handers in the left (*t*(190.1) = 3.026, *p* = 0.012, *n* = 4) but not in the right hemisphere (*t*(190.1) = −1.624, *p* = 0.424, *n* = 4).
– Orthogonal: A significant main effect of the hemisphere (*F*(1,96) = 11.283, *p* = 0.001, 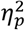 = 0.105) was shown so that the absolute orthogonal coordinate was larger in the left than in the right hemisphere. A significant effect of the interaction between sex and handedness categor*F*y((1,96) = 6.782, *p* = 0.011, 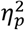 = 0.066) was also reported. For males, Low right-handers had a larger absolute orthogonal coordinate than the High right-handers (*t*(96) = −3.390, *p* = 0.004, *n* = 4) which was not the case for females (*t*(96) = 0.210, *p* = 1, *n* = 4). Moreover, no significant difference was found between males and females either in the High right-handers (*t*(96) = 1.618, *p* = 0.436, *n* = 4) or in the Low right-handers (*t*(96) = −2.113, *p* = 0.148, *n* = 4).
- Centroid 4:

– Diagonal: A significant interaction between hemisphere and handedness category was assessed (*F*(1,96) = 16.935, *p* < 0.001, 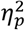 = 0.119). Centroid was more dorsal in the left than in the right hemisphere in the High right-handers (*t*(96) = 4.147, *p* < 0.001, *n* = 4) but not in the Low right-handers (*t*(96) = −0.694, *p* = 1, *n* = 4). Moreover, centroid was more dorsal in the High right-handers than in the Low right-handers in the left (*t*(186,8) = 3.058), *p* = 0.012, *n* = 4) but not in the right hemisphere (*t*(186,8) = −1.586, *p* = 0.456, *n* = 4).
– Orthogonal: No significant effect of either handedness category, sex or hemisphere was found on this coordinate.
- Centroid 5: No significant effect of any of the factors on either diagonal or orthogonal coordinate of centroid 5 was reported.

The position of all centroids, with the notable exception of centroid 5 at the dorsal end of the CS, therefore, depends on hemisphere, sex and manual laterality score. In addition, for the centroids 2, 3 and 4 located around the hand area, there is an interhemispheric asymmetry of position along the axis of the CS (*Diag_C* coordinate) that depends on the manual laterality score. This asymmetry reveals a more dorsal position of centroids 2, 3 and 4 in the left hemisphere for subjects with high manual laterality scores. For these same bundles, the effect sizes associated with the results of the *Orth_C* coordinate underline that the value of the latter depends essentially on the hemisphere considered.

### Streamlines count and volume of the bundles

A significant leftward asymmetry of streamlines count was shown in bundle 1 (*t*(99) = −3,393, *p* < 0.001), bundle 3 (*t*(99) = −6.621, *p* < 0.001) and bundle 5 (*W* = 1203, *p* < 0.001) and this asymmetry was not dependent on sex or handedness category (all p>0.05) for bundles 1 and 3. However, a significant effect of handedness category (*U* = 921, *p* = 0.025) but not of gender (*U* = 1052, *p* = 0.148) on streamlines count asymmetry was assessed for bundle 5. Regarding bundle’s volume, a leftward asymmetry was reported for bundle 1 (*W* = 1256, *p* < 0.001) and bundle 2 (*t*(99) = −8.00, *p* < 0.001).

A linear regression was calculated to predict dexterity score based on streamlines count of each three of the central clusters of the left hemisphere covarying for sex. No significant regression equation was found for any of the clusters (all *p* ≥ 0.05). Similarly, a linear regression was used to predict strength score with respect to streamlines count asymmetry of each three central clusters, covarying for sex but here also no significant regression equation was found for any of the clusters (all *p* ≥ 0.05).

## Discussion

In this study we proposed the definition of the connectivity space, a two-dimensional cartesian domain, parameterized by the gyral crests adjacent to the sulcus. In this space, a short association streamline coursing around the sulcus is defined as a point whose coordinates are the positions of its end points on each gyral crest. Compared to the manual seed placement bundle selection approach used notably in (Catani et al. 2012; Rojkova et al. 2016), the connectivity space allowed for a more comprehensive and continuous description of U-shape fibres structural connectivity around a sulcus without seed-related prior. This connectivity space has been used to construct individual and group connectivity profiles of the left and right CS from the 100 subjects.

The concentration of fibre terminations close to the diagonal of the group connectivity spaces as well as the presence of density maxima of these terminations along the dorso-ventral axis and in functionally distinct zones are in favour of a functional organization of U-shape fibre bundles, i.e., U-shape fibres mainly connecting functionally homologous territories of the primary somatosensory area as suggested in (Catani et al. 2012) from delineated discrete bundles. In a recent study (Germann et al. 2019), showed a close anatomo-functional correspondence between prominences of the walls, plis de passage and the functional areas of the CS. Our results also suggest, as conjectured in (Guevara et al. 2018), an additional correspondence between these morphological features of the CS and U-fibres terminations as evidenced by the presence at the group level of U-shape fibre density maxima at the PPFM location.

When constructing a discrete representation from continuous group connectivity profiles, five bundles were identified bilaterally: one bundle in the functional area of the tongue (ventral end), three bundles in the functional area of the hand and one in the functional area of the foot (dorsal end). If the overall organisation and number of bundles we obtained along the CS is in line with the literature (e.g. (Catani et al. 2012; Rojkova et al. 2016; Guevara et al. 2017b, 2018; Román et al. 2017)), especially in the functional hand area, some differences can be noted, however. Hence, in the ventral end, Catani et al. (2012) distinguished two bundles, namely *the lower and upper face bundles*. Nevertheless, (Guevara et al. 2017; Román et al. 2017) also obtained a single stable bundle in the ventral end of the CS by clustering the streamlines based of the whole streamline geometry. This implies that the obtaining of a single bundle cannot be attributed solely to the consideration of the termination of the fibres in defining the bundles. Visual inspection of the individual U-shape streamlines we obtained revealed a possible subdivision of the ventral bundle into two sub-bundles in some subjects but not in all. Guevara et al. (2017b) studied the stable bundles of the CS with respect to homogeneous morphological configurations of the CS and also exhibited configurations with two fibre bundles and configurations with a single fibre bundle in the ventral end. It is therefore likely that our single ventral bundle finding could actually be a group effect hiding two different subgroups. Concerning the dorsal end of the CS, a bundle was obtained corresponding to the *paracentral bundle* extracted in (Catani et al. 2012). Hence again, the visual inspection of this bundle revealed it could be subdivided into at least two sub-bundles in some subjects. These reported differences could mostly be attributed to inter-individual variability in the number and position of the bundles. Such variability sometimes interferes with the identification of some bundles, depending on the quality of the data and the algorithms. For instance, Thompson et al., (2017) were not able to extract consistently the paracentral tract from dMRI data with 2.4 mm isotropic resolution, 32 diffusion-weighted directions and tractography based on the diffusion tensor model with no consideration of the partial volume effect. The variability is likely to be critical in our study since our tractograms include short streamlines (between 3 and 20 mm), that are usually not considered especially in clustering-based studies (e.g. (Guevara et al. 2017; Román et al. 2017)).

We have then quantitatively shown that the centroids of the bundles of the left CS hand area of the strongly right-handed subjects are more dorsal than those of the weakly right-handed and those of the right hemisphere. Our results confirm the qualitatively reported in (Guevara et al. 2018), dorsal *translation* of the ends of the bundles of the hand area according to the variation in the morphology of the CS which had been previously linked with handedness (Sun et al. 2016). This dorsal translation of the motor (and sensory) functions of the hand was associated with the lateralization of language functions in the left hemisphere in right-handed people (Karolis et al. 2019), including a transfer of the motor regions of the larynx and lips, from the walls of the pre-central sulcus to a bulge of the pre-central gyrus ventrally located in the handknob (Sun et al. 2016). A secondary implication of our result is that the manual laterality score could be more closely associated than what was previously done (left and right handers distinction) in (Sun et al. 2012) with the variability of the morphology of the CS and its short structural connectivity.

Interhemispheric asymmetries in the volume of the bundles and in the number of streamlines making up these bundles in the functional area of the hand do not depend on any functional or phenotypic score tested in this study and therefore appears to be *structural*. For, the central bundle of the hand area, the greater number of streamlines in the left than in the right hemisphere and the absence of volume difference suggests a higher density of the streamlines making this bundle in the left hemisphere. Conversely, the larger volume of the central lower bundle coupled with the identical number of streamlines in this bundle suggests a larger spatial dispersion of the bundle and/or a longer average bundle. Hence, at most it implies that geometric and microstructural properties of bundles are different between hemispheres. Our results regarding volume and streamlines count asymmetries are in line with those of (Catani et al. 2012) but in opposition to those of (Magro et al. 2012; Román et al. 2017) which report respectively an absence of asymmetry in the number of streamlines for right-handed subjects and an asymmetry in favour of the right hemisphere of the volume of the lower and central bundles of the hand region. Direct comparison of these results is complex, however. Indeed, the number of streamlines or the voxel volume of a bundle does not necessarily reflect an underlying biological reality but rather methodological confounds such as the initialization strategy of the tractography used as well as the acquisition artefacts (Jones et al. 2013; Smith et al. 2015b). Unlike previously referenced studies, we have tried as much as possible to limit the influence of these confounds, by using state-of-the-art and robust preprocessing pipeline including correcting the data for antenna bias and filtering the tractograms obtained with respect to the dMRI signal (Daducci et al. 2015).

Regarding the CS U-fibres functional role, studies carried out on animal models, e.g. rodents, monkeys, that also have direct connections between primary somatosensory areas, have shown a predominant role of local extrinsic connections from the primary sensitive zone to the primary motor zone in motor learning (see (Papale and Hooks 2018) for a review) and fine grasping (Hikosaka et al. 1985). Thompson et al., (2017) reported a correlation in humans between diffusion tensor metric values for bundles in the functional area of the left hand and manual dexterity scores in both hands suggesting a dominant role of left CS U-fibres in fine motor control. In this study, we did not observe a correlation between the number of streamlines in the left hemisphere and the fine manual dexterity score, a result that is also consistent with the absence of correlation between the asymmetry of the streamline number in the central bundle and manual laterality, manual dexterity and manual laterality being linked. Also, given that the metrics used in (Thompson et al. 2017) were defined at the voxel level, there is no explicit identification of fibre bundles and the correlation reported could be associated to other bundles such as thalamocortical projections or corticospinal tracts. If the asymmetry of the number of streamlines is not related to manual dexterity, it could then be involved in motor learning: asymmetry is present in the tongue and hand areas, two parts of the body involved in functions requiring complex motor learning e.g. fine grip, language. The putative involvement of these fibres in motor learning is consistent with the temporality of the myelination of these fibres which occurs up to the approximate age of two years (Dubois et al. 2014). However, this presupposes a decorrelation of motor learning and manual performance.

## Limitations

While dMRI and associated reconstruction techniques (local modelling, tractography) have led to considerable advances in knowledge of the macro- and micro-structure of human white matter (Assaf et al. 2019), they are not free of bias (Maier-Hein et al. 2017; Schilling et al. 2018, 2019) that can affect neuroscientific conclusions (Sinke et al. 2018). In this study, we tried to reduce the impact of already identified bias by relying on high quality pre-processed data available in the HCP dataset, taking into account partial volume effect in the local modelling step, adopting an anatomically constrained probabilistic tracking algorithm capable of estimating highly curved fibres while removing streamlines stopping in the white matter and filtering the tractogram using a state-of-art technique (Daducci et al. 2015) in order to obtain tractograms representative of the dMRI signal and thus more quantitative streamlines counts. Despite these methodological precautions, results obtained should be treated with caution, particularly since estimated streamlines are organized in complex configurations (Reveley et al. 2015), and since results may dependent on pre-processing pipeline (e.g. (Maier-Hein et al. 2017; Brun et al. 2019; Maximov et al. 2019)).

In addition, the connectivity space framework presents some limitations. This connectivity space is built using streamline endpoints whose position is likely to be affected by noise and partial volume effect. In addition, since this framework relies on a projection on the crests of the adjacent gyri, it implies some knowledge is lost in other dimensions. As an example, (Viganò et al. 2019) reported a rostro-caudal functional gradient (perpendicular to the sulcus) in the hand area an axis along which the structural connectivity is averaged in our study.

Finally, the bundle selection process and the inter-subject correspondence could be improved in order to better account for inter-subject morphological variability. In this study, we chose to only align the PPFM position across subjects using piecewise affine transformation as a compromise between deformation of the initial space and the alignment accuracy. However, more complex non-linear registration methods could be used in order to account for more subtle variations of the anatomy. Regarding streamlines bundle extraction, a two-step clustering approach such as in (Guevara et al. 2017a; Román et al. 2017) may improve the inter-subject correspondence by disentangling cases where homologous bundles have different position around the CS. It could be used in conjunction with advanced correspondence technics such as optimal transport to map more accurately structural connectivity profiles across individuals.

## Conclusion

Based on the anatomical connectivity space, which allows an exhaustive and continuous representation of the anatomical connectivity of the U-shape fibres of a sulcus, our study corroborated a functional organization of the latter in the case of the human CS, and highlighted a difference in the position of the hand bundles of the contralateral hemisphere in relation to handedness. Our findings thus suggest an important correspondence, between the CS morphology, short structural connectivity and manual preference. A direct extension of this work would involve processing the entire HCP S1200 release dataset in order to further characterize the U-fibres of the CS and particularly better quantify the effect of handedness (right versus left handers) on the short U fibre connectivity of the CS. Another complementary line of research would be to characterise the position of the endings of these fibres in relation to the morphological landmarks of a sulcus such as its geometric bulges or its *pli de passage* at both the individual and the group level. The connectivity space framework could also be applied to other sulci of the human brain such as the superior temporal sulcus, in conjunction with detailed functional characterization in order to try characterizing it’s U fibres structural variability and at least some of the functional roles it’s U-fibres may have in a precise context.

## Acknowledgements

The authors would like to thank Pr. Alessandro Daducci for his precious advice and discussion about the COMMIT framework.

Data were provided by the Human Connectome Project, WU-Minn Consortium (Principal Investigators: David Van Essen and Kamil Ugurbil; 1U54MH091657) funded by the 16 NIH Institutes and Centers that support the NIH Blueprint for Neuroscience Research; and by the McDonnell Center for Systems Neuroscience at Washington University.

## Declarations

### Funding

Alexandre Pron is supported by doctoral a grant from Aix-Marseille University.

### Conflicts of interest

The authors declare that they have no conflict of interest.

### Code availability

The code used to carry out the analyses of the current study is publicly available at https://github.com/alexpron/article_central_sulcus_connectivity

### Authors’ contributions

All authors contributed to the study conception and design. Alexandre Pron implemented the processing pipeline and performed meshes visual quality control. Olivier Coulon and Alexandre Pron manually drew the surface cortical landmarks of the central sulcus onto the grey matter/white matter interface meshes. All authors contributed to the statistical analyses, the preparation, the writing, and the correction of the article. All authors read and approved the final manuscript.

## Supplementary Material

### CS endpoints positioning

The PreCG, PoCG, and CS fundus lines were imposed to have the same endpoints. Dorsal endpoint was set at the apex of the characteristic notch made by the CS on the medial face of the GM/WM mesh. Ventral endpoint was set on the crest of the sub-central gyrus (SubCG). Precise positioning of the endpoints was achieved using the Depth Potential Function (DPF) (Boucher et al. 2009), a regularized scalar field which includes both mean curvature and average convexity information. An endpoint was set at the vertex of an extremal area (e.g. the SubCG) that was closest to the 0-level set of the DPF and whose value was negative (see **Fig. *6***).

In contrast to the paracentral lobule, the ventral part of the CS exhibited significant morphological and topological variability as illustrated in **Fig. *6***. Anatomical descriptions from (Duvernoy et al. 1999; Scarabino 2006; Petrides 2019) were used to consistently locate the SubCG. First, main gyri and sulci of the central area (i.e., the PreCG, the PoCG, the pre-central sulcus (PreCS) and the post-central sulcus (PoCS)) were identified on the pial mesh as shown **Fig. *6***. We then sought the anterior sub-central sulcus (a-SubCS) and the posterior sub-central sulcus (p-SubCS) as two notches on the superior border of the median lateral fissure. More precisely, the a-SubCS notch started between the inferior PreCS and the PreCG, whereas the p-SubCS notch was between the inferior PoCG and the PoCS. In some subjects, at least one notch was not directly visible on the pial mesh but was visible on the GM/WM mesh. The SubCG was pinpointed between the a-SubCS and the p-SubCS. In addition, we used the posterior branch of the PreCS (p-PreCS), when recognizable, as a complementary criterion to disentangle configurations with two interruptions: moving ventrally from the intersection between the p-PreCS and the CS, the SubCG first interrupted the CS.

### PreCG and PoCG crest lines extraction

The PreCG and PoCG crests were extracted as the shortest path between the two extremal point weighted by the DPF. Four manually specified controls points were added to differentiate between gyri and to ensure an accurate delineation in bending areas. Controls points of the PreCG were placed from the ventral to the dorsal extremity at: 1) the intersection with the p-PreCS, 2) the ventral extremity of the first genu of the hand-knob area, 3) the dorsal extremity of the last genu of the hand-knob area and 4) the intersection with the superior frontal gyrus. PoCG control points were set at major bending locations and were usually counterpart of the ones of the PreCG as shown.

### PPFM delineation

The position of the PPFM along the CS fundus line was set manually. The *hand-knob*, a characteristic backward bulge (Yousry et al. 1997) often located near the midpoint of the CS was first identified on the GM/WM mesh. In the case of *double knob* configuration, as described in (Sun et al. 2012), the knob with the thicker protrusion in the CS was preferred. The PreCG backward hand-knob protrusion faced another one, coming forward from the PoCG, and often more ventral, leading to an interdigitating pattern (e.g. (White 1997)). The PPFM location was set at the crossing between the CS line and a line joining both extremities of the protrusion in the sulcus fundus where a slight and focal elevation of the sulcal floor was usually observed (Boling and Reutens 1999).

In very rare cases where the PPFM explicitly split the CS into two components (2 hemispheres out of 200), its location was set at the intersection of the sulcal line and the crest of the pli as shown on the bottom line of **Fig. *7***.

PPFM of the 100 subjects were located by two operators (O.C. or A.P.). No statistically significant difference in position between operators was reported in either hemisphere (Mann-Whitney tests between operators positions, one for each hemisphere, *U*_1_ = 747, *p*_1_ = 0.24, *U*_*r*_ = 632, *p*_*r*_ = 0.17). PPFM projections on both adjacent gyral crests lines were obtained by minimizing the geodesic distance to these lines weighted by mean curvature (Shattuck et al. 2009). The mean curvature constraint reduced the variability induced by the mesh geometry in this area and consistently projected the PPFM on the middle of the bulge of the knob and on the corresponding bend of the PreCG as shown **Fig. *7***.

## Footnotes

1 https://www.humanconnectome.org/study/hcp-young-adult

## Notes

https://github.com/alexpron/article_central_sulcus_connectivity

### Competing Interest Statement

The authors have declared no competing interest.

